# LEF1-AS1 deregulation in the peripheral blood of patients with persistent post-COVID symptoms

**DOI:** 10.1101/2024.12.20.629131

**Authors:** Alisia Madè, Santiago Nicolas Piella, Marco Ranucci, Carlo Gaetano, Laura Valentina Renna, Rosanna Cardani, Gaia Spinetti, Fabio Martelli

## Abstract

Long COVID denotes the persistence of symptoms after acute SARS-CoV-2 infection lasting for at least two months without another identifiable cause. Affecting an estimated 15% of COVID-19 patients, long COVID manifests in a wide range of symptoms. Despite extensive research on its one-year effects, limited data exist beyond 12 months. Due to the different manifestations of long COVID, its diagnosis can be challenging. Identifying potential mechanistic contributors and biomarkers would be highly valuable. Recent studies have highlighted the potential of noncoding RNAs (ncRNAs) as biomarkers for disease stratification in COVID-19. Specifically, we have recently identified miR-144-3p and a subset of lncRNAs as candidates for assessing disease severity and outcomes in COVID-19. This study extends such investigations to 98 long COVID patients recruited 18 months after hospitalization, exploring the relationship between circulating ncRNA expression and persistent symptoms. While miR-144-3p, HCG18, and lncCEACAM21 expression did not differ between symptomatic and asymptomatic patients, LEF1-AS1 was downregulated in peripheral blood mononuclear cells (PBMCs) of symptomatic patients. Of note, multiple LEF1-AS1 isoforms and LEF1 sense transcript levels were reduced and negatively correlated with relevant clinical markers. While further studies are needed, our discoveries offer new perspectives for the diagnosis and management of persistent long COVID.

## 1. Introduction

Long COVID, also known as post-COVID, is a multisystem condition defined as the continuation or emergence of new symptoms three months after SARS-CoV-2 infection, persisting for at least two months without any other identifiable cause [1]. The World Health Organization (WHO) has estimated that 10-20% of patients who suffered from COVID-19 develop long COVID symptoms [2]. Although some studies suggest that COVID-19 vaccination reduces the risk of developing long COVID symptoms [3–5], it remains a significant global issue [6]. This challenge is partly due to unequal access to vaccines and booster doses. Despite some studies indicate that the severity of COVID-19 infection may have an impact on the probability to develop long COVID symptoms [7,8], a European Respiratory Society statement reported that the risk factors for long COVID are not necessarily associated with the initial severity of the illness. Instead, they could be linked to female sex, age, and the number of symptoms experienced at onset [9,10]. Common symptoms of long COVID include fatigue, shortness of breath, and cognitive impairment, but the condition is highly variable, affecting numerous organs [11,12]. The pathogenesis of long COVID is not yet fully understood, but it is thought to involve a combination of persistent viral infection, immune dysregulation, and residual tissue damage. These factors may contribute to the diverse range of symptoms experienced by patients [11].

Although numerous studies have investigated the long-term effects of COVID-19 one year after infection [13] mainly focusing on pulmonary functions and respiratory outcomes [14–16], and cognitive deficits [17], there are limited data on the condition beyond 12 months from the onset of COVID-19 [5,18–21].

Due to the different manifestations of long COVID, the diagnosis of the syndrome can be challenging. Hence, identifying potential mechanistic contributors and biomarkers would be highly valuable. In line with this evidence, noncoding RNAs (ncRNAs) have shown great promise as biomarkers for various diseases, including COVID-19. ncRNAs can be classified as small noncoding RNAs (e.g. microRNA) or long noncoding RNAs (lncRNAs), based on their transcript size. ncRNAs can regulate the transcription and translation of protein-coding RNAs [22] and are also detectable in peripheral blood, providing potential as disease biomarkers.

In previous studies investigating acute patients, we explored the expression levels of a specific microRNA (miRNA), miR-144-3p, in plasma or serum samples from three different groups of COVID-19 patients recruited across Europe, including both hospitalized and non-hospitalized individuals. qPCR analysis identified miR-144-3p as a promising candidate for risk-based stratification and mortality prediction in COVID-19 illness [23]. Among the lncRNAs, HCG18-family lncRNAs, lncCEACAM21, and LEF1-AS1-202 lncRNAs are downregulated in PBMC of non-surviving COVID-19 patients compared to surviving ones, as well as in critical compared to severe patients, thus emerging as potential candidates for discriminating severity and predicting mortality rates in hospitalized COVID-19 patients [24]. Accordingly, a machine-learning model based on age and lncRNA LEF1-AS1 can predict the outcome of hospitalized COVID-19 patients in large groups of European and Canadian patients [25].

In this study, we investigated the expression of selected COVID-19 ncRNAs in a cohort of 98 long COVID patients enrolled 18 months from COVID-19 hospitalization and assessed their potential role as biomarkers of the presence of long COVID symptoms.

## 2. Results

### 2.1 Long COVID patient characteristics

For this study, 98 patients hospitalized at IRCCS Policlinico San Donato because of mild to severe COVID-19 were enrolled in the “long COVID” protocol to assess symptoms persistent after hospitalization. Table 1 summarizes the general characteristics of the patients during their acute phase, including COVID-19 pre-existing conditions and information about in-hospital treatments.

**Table 1.**
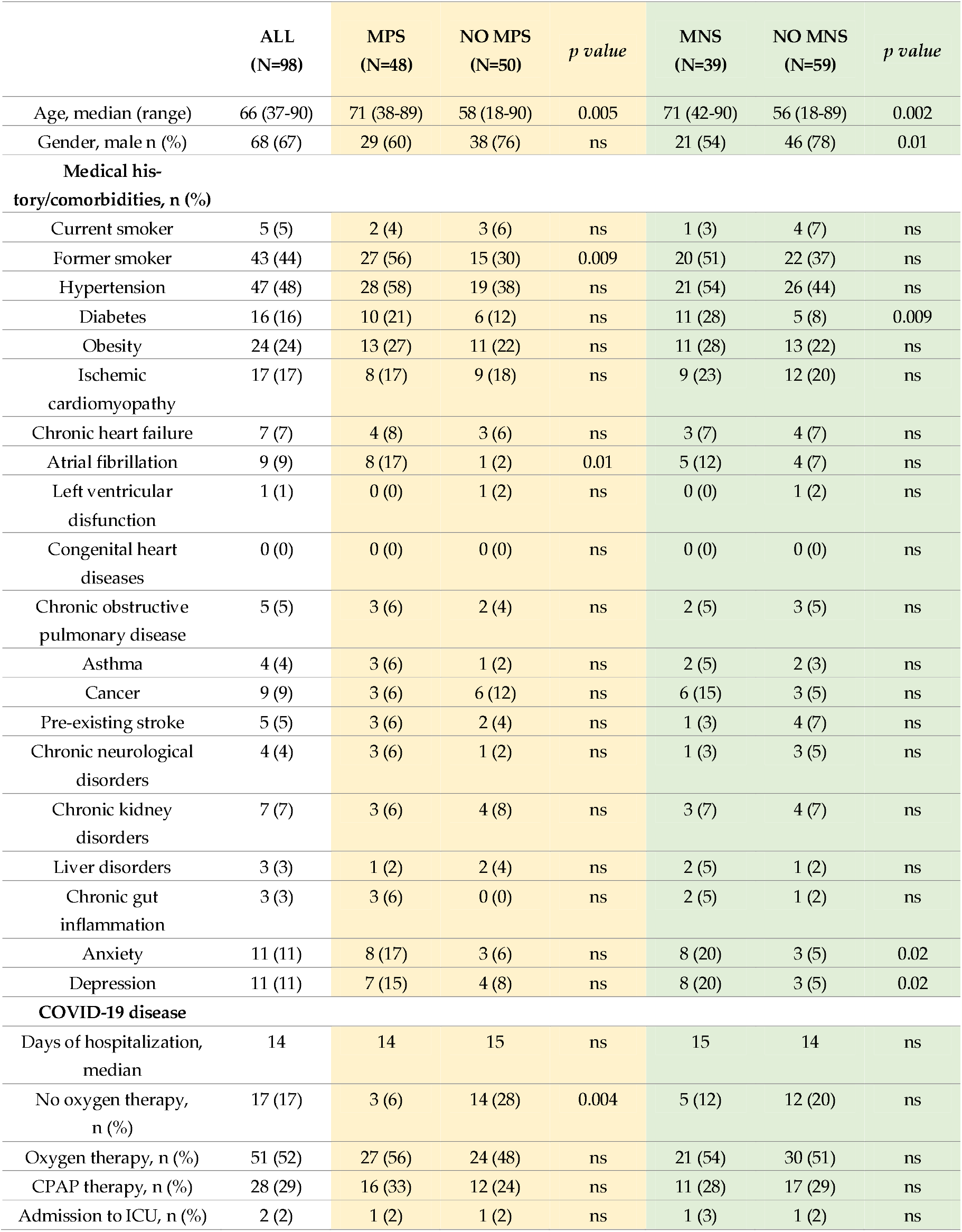
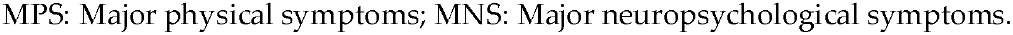
Patient characteristics at hospital admission and during acute COVID-19 disease.

|  | ALL<br>(N=98) | MPS<br>(N=48) | NO MPS<br>(N=50) | *p value* | MNS<br>(N=39) | NO MNS<br>(N=59) | *p value* |
| --- | --- | --- | --- | --- | --- | --- | --- |
| Age, median (range) | 66 (37-90) | 71 (38-89) | 58 (18-90) | 0.005 | 71 (42-90) | 56 (18-89) | 0.002 |
| Gender, male n (%) | 68 (67) | 29 (60) | 38 (76) | ns | 21 (54) | 46 (78) | 0.01 |
| **Medical history/comorbidities, n (%)** |  |  |  |  |  |  |  |
| Current smoker | 5 (5) | 2 (4) | 3 (6) | ns | 1 (3) | 4 (7) | ns |
| Former smoker | 43 (44) | 27 (56) | 15 (30) | 0.009 | 20 (51) | 22 (37) | ns |
| Hypertension | 47 (48) | 28 (58) | 19 (38) | ns | 21 (54) | 26 (44) | ns |
| Diabetes | 16 (16) | 10 (21) | 6 (12) | ns | 11 (28) | 5 (8) | 0.009 |
| Obesity | 24 (24) | 13 (27) | 11 (22) | ns | 11 (28) | 13 (22) | ns |
| Ischemic cardiomyopathy | 17 (17) | 8 (17) | 9 (18) | ns | 9 (23) | 12 (20) | ns |
| Chronic heart failure | 7 (7) | 4 (8) | 3 (6) | ns | 3 (7) | 4 (7) | ns |
| Atrial fibrillation | 9 (9) | 8 (17) | 1 (2) | 0.01 | 5 (12) | 4 (7) | ns |
| Left ventricular disfunction | 1 (1) | 0 (0) | 1 (2) | ns | 0 (0) | 1 (2) | ns |
| Congenital heart diseases | 0 (0) | 0 (0) | 0 (0) | ns | 0 (0) | 0 (0) | ns |
| Chronic obstructive pulmonary disease | 5 (5) | 3 (6) | 2 (4) | ns | 2 (5) | 3 (5) | ns |
| Asthma | 4 (4) | 3 (6) | 1 (2) | ns | 2 (5) | 2 (3) | ns |
| Cancer | 9 (9) | 3 (6) | 6 (12) | ns | 6 (15) | 3 (5) | ns |
| Pre-existing stroke | 5 (5) | 3 (6) | 2 (4) | ns | 1 (3) | 4 (7) | ns |
| Chronic neurological disorders | 4 (4) | 3 (6) | 1 (2) | ns | 1 (3) | 3 (5) | ns |
| Chronic kidney disorders | 7 (7) | 3 (6) | 4 (8) | ns | 3 (7) | 4 (7) | ns |
| Liver disorders | 3 (3) | 1 (2) | 2 (4) | ns | 2 (5) | 1 (2) | ns |
| Chronic gut inflammation | 3 (3) | 3 (6) | 0 (0) | ns | 2 (5) | 1 (2) | ns |
| Anxiety | 11 (11) | 8 (17) | 3 (6) | ns | 8 (20) | 3 (5) | 0.02 |
| Depression | 11 (11) | 7 (15) | 4 (8) | ns | 8 (20) | 3 (5) | 0.02 |
| **COVID-19 disease** |  |  |  |  |  |  |  |
| Days of hospitalization, median | 14 | 14 | 15 | ns | 15 | 14 | ns |
| No oxygen therapy, n (%) | 17 (17) | 3 (6) | 14 (28) | 0.004 | 5 (12) | 12 (20) | ns |
| Oxygen therapy, n (%) | 51 (52) | 27 (56) | 24 (48) | ns | 21 (54) | 30 (51) | ns |
| CPAP therapy, n (%) | 28 (29) | 16 (33) | 12 (24) | ns | 11 (28) | 17 (29) | ns |
| Admission to ICU, n (%) | 2 (2) | 1 (2) | 1 (2) | ns | 1 (3) | 1 (2) | ns |

MPS: Major physical symptoms; MNS: Major neuropsychological symptoms.

In agreement with a previous study performed at our hospital analysing a similar patient group, long COVID patients were classified for the presence or absence of major physical symptoms (MPS) or major neuropsychological symptoms (MNS) [5], as assessed at follow-up. Only 2% of the entire population required admission to the Intensive Care Unit (ICU), while the majority required oxygen therapy during their stay in the hospital. After a median of 18 months from the onset of COVID-19, patients were enrolled in the long COVID protocol and asked about the presence or development after discharge of symptoms recorded in Table 2.

**Table 2.**
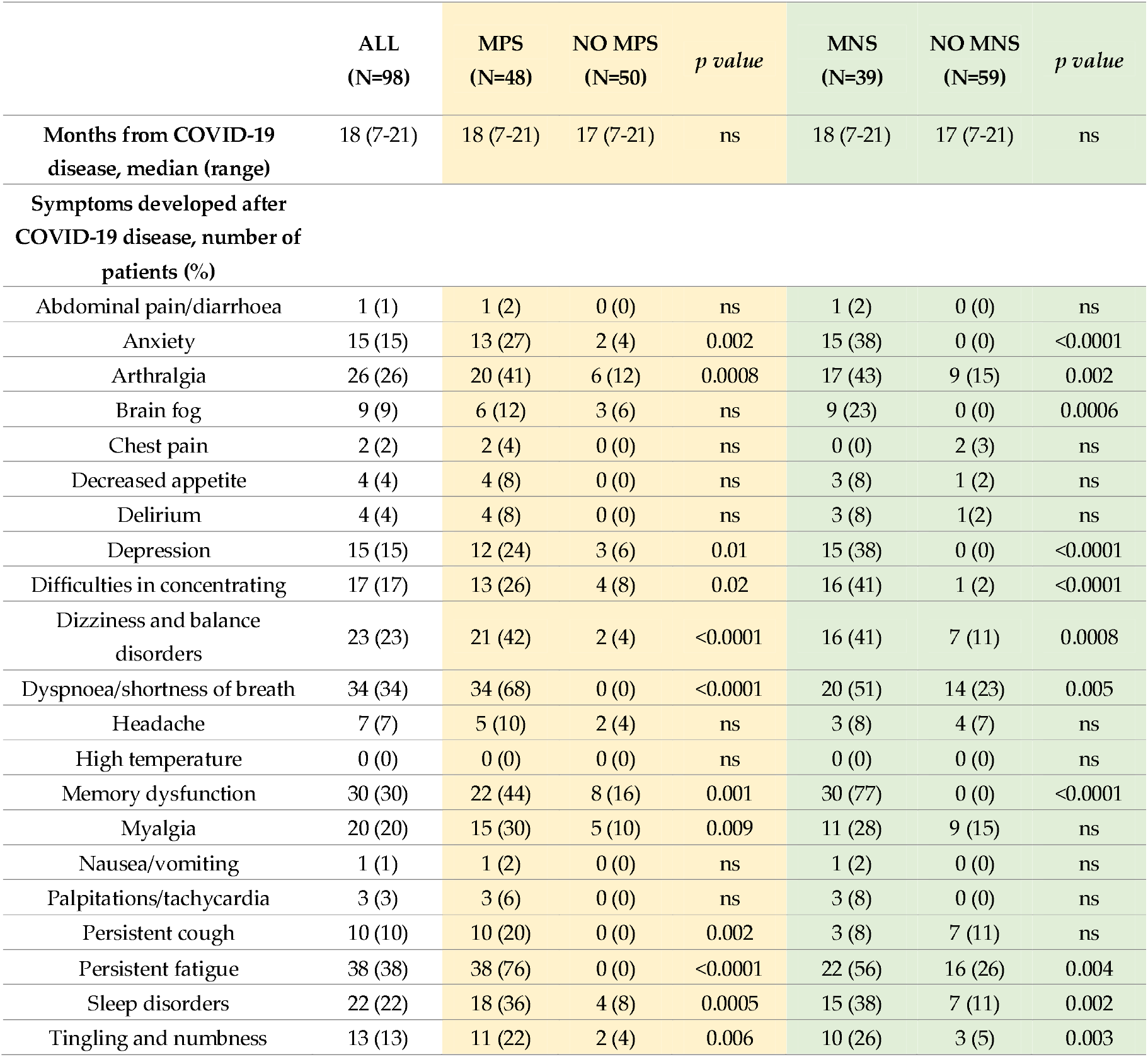
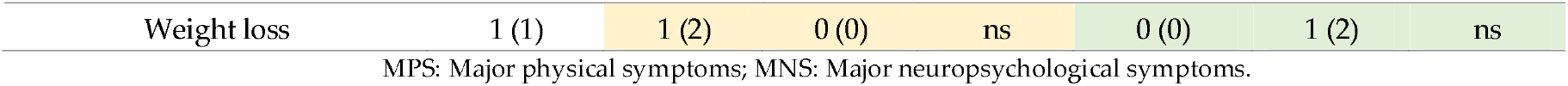
Symptoms developed after hospital discharge and are still present at the moment of the interview.

Among symptoms still present at the time of follow-up, the most frequent ones were persistent fatigue (38%), dyspnoea (34%) and memory dysfunction (30%). MPS were more frequent in former smokers and, conversely, less frequent in those patients not requiring oxygen therapy during the hospital stay, possibly indicating a milder form of the acute disease (Table 1). Persistent MNS were more frequent in patients that were already diabetics at the time of acute COVID-19 (Table 1).

### 2.2 No difference in plasma miR-144-3p levels in MPS or MNS long COVID patients

To investigate whether the development or the absence of long COVID symptoms was associated with persistent dysregulation of plasmatic miR-144-3p [23], its expression levels were measured in platelet-poor plasma samples of the long COVID patients. Results revealed no significant differences in miR-144-3p expression levels in the comparison of patients with or without MPS or MNS (Figure S1).

### 2.3 Lower LEF1-AS1 expression levels in the PBMC of MPS patients

To determine whether COVID-19 lncRNA biomarkers [24] remain persistently dysregulated, potentially indicating the development of long COVID symptoms, RNA extracted from PBMC samples of long COVID patients was analysed. LEF1-AS1-202 displayed significantly decreased levels in MPS patients compared to those without MPS (Figure 1A), but not in patients affected by persistent MNS compared to their controls (Figure S2A). Conversely, the expression of HCG18 (both HCG18-244 and long isoforms) and lncCEACAM21 displayed no significant differences between patients with or without MPS (Figure 1 B, C and D), or MNS (Figure S2B, C, and D). Thus, LEF1-AS1 expression in the PBMCs of long COVID patients was further explored.

**Figure 1.**
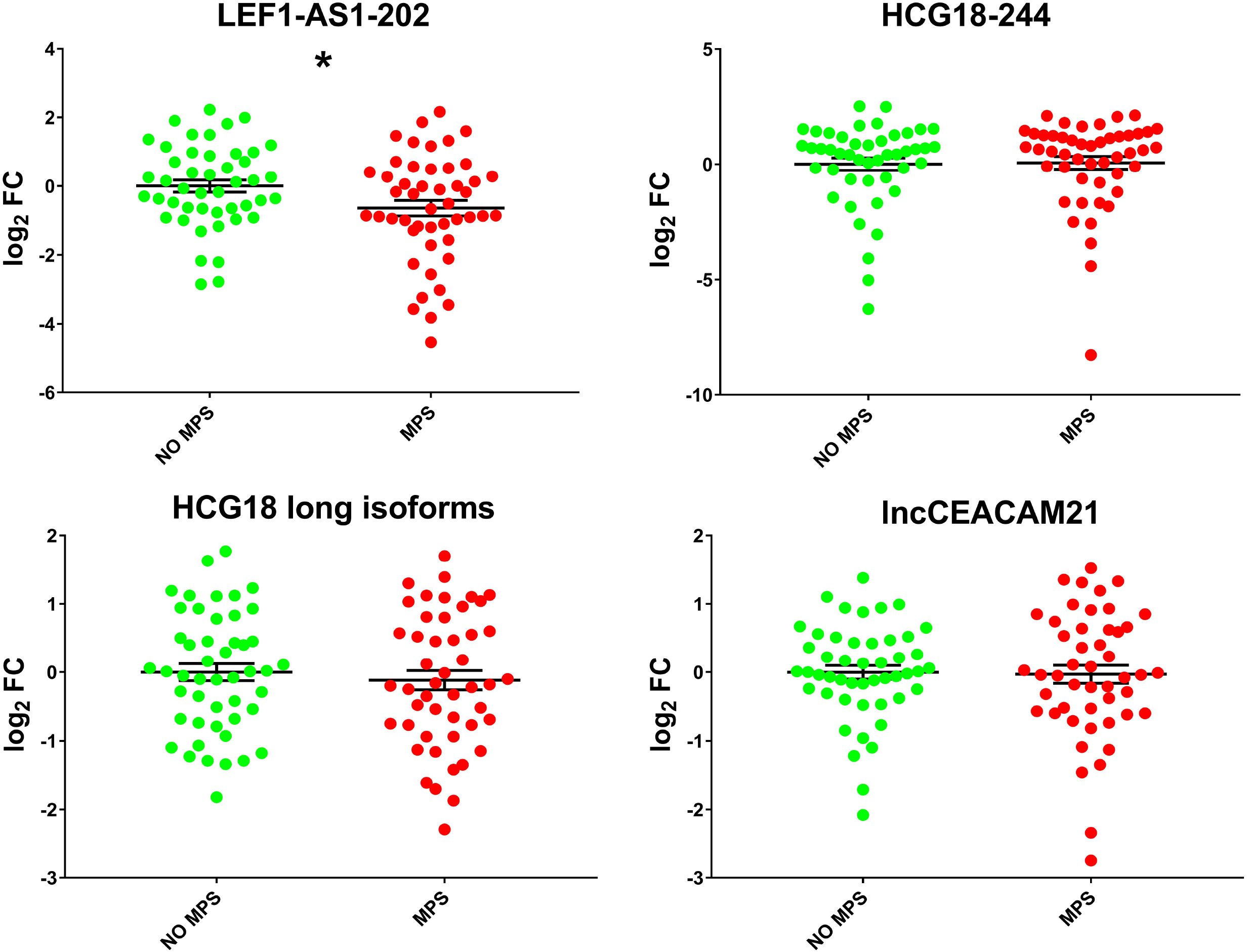
Lower LEF1-AS1-202 levels in MPS patients compared to no MPS patients. The levels of LEF1-AS1-202 (A) HCG18-244 (B) HCG18 long isoforms (C) and lncCEACAM21 (D) were measured in RNA extracted from PBMCs derived from patients with (MPS, n = 48) or without MPS (no MPS, n = 46-48). LEF1-AS1-202 (A) showed a significant decrease in MPS patients compared to no MPS patients (* p < 0.05), while no significant differences were observed in the expression levels of the HCG18-244 (B), HCG18 long isoforms (C) and lncCEACAM21 (D). Values are expressed as log2 fold change and shown as dot-plots, indicating mean ± SEM. Unpaired t-test (two groups) was used for statistical comparison.

### 2.4 LEF1-AS1 isoforms and LEF1 are similarly deregulated in PBMCs of long COVID patients

Different LEF1-AS1 isoforms have been annotated in GRCh38.p14 (Figure S3). In line with LEF1-AS1-202 modulation, other LEF1-AS1 isoforms were investigated in long COVID patients. Specific primers were designed to detect LEF1-AS1 –210, –211, –204 –212, and –209 isoforms collectively (referred to as LEF1-AS1 multiple isoforms), or LEF1-AS1-207 isoform. Additionally, to validate the possible co-regulation with LEF1, its expression levels were also analysed. Decreased levels of all LEF1-AS1 isoforms tested were observed in patients affected by MPS compared to their no-MPS controls (Figure 2A-B), consistent with the LEF1-AS1-202 result. Furthermore, LEF1 expression levels were also lower in patients with MPS compared to those without MPS (Figure 2C), indicating a similar regulation of LEF1-AS1 and LEF1.

**Figure 2.**
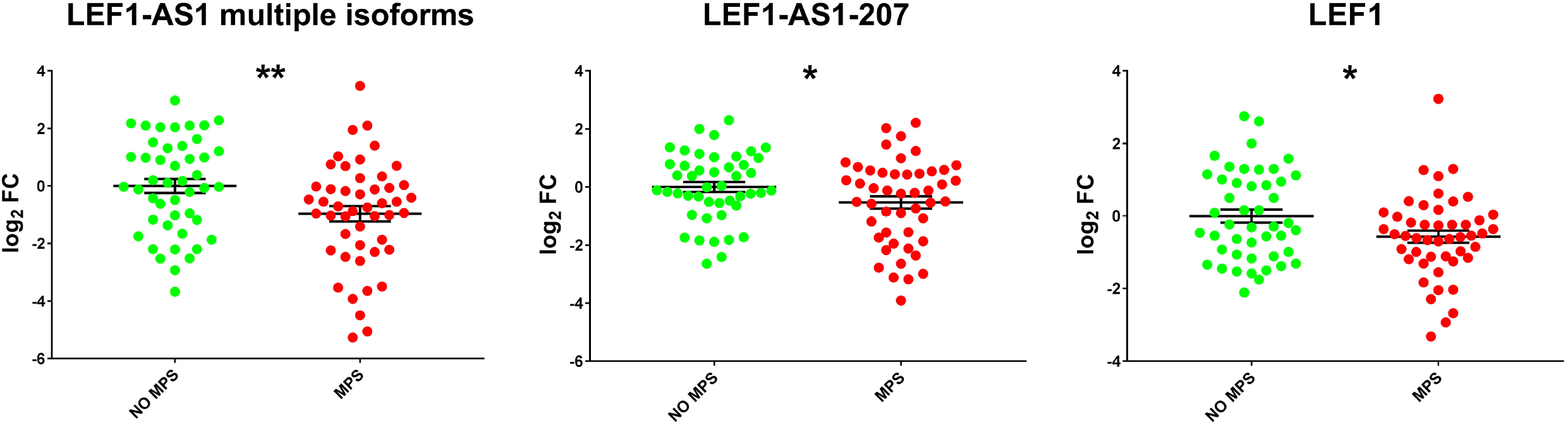
Lower levels of LEF1-AS1 isoforms and LEF1 in MPS patients. LEF1-AS1 multiple isoforms (A), LEF1-AS1-207 (B), and LEF1 (C) were measured in total RNA extracted from the PBMCs of patients with (MPS, n = 48) or without MPS (no MPS, n = 46). Values are expressed as log2 fold change and shown as dot-plots, indicating mean ± SEM. Unpaired t-test (two groups) was used for statistical comparison. * p < 0.05; ** p ≤ 0.01.

Accordingly, LEF1 levels correlated positively to the expression of LEF1-AS1-202 (r(S) = 0.71), of LEF1-AS1 multiple isoforms (r(S) = 0.79) and of LEF1-AS1-207 (r(S) = 0.63) (Figure 3).

**Figure 3.**
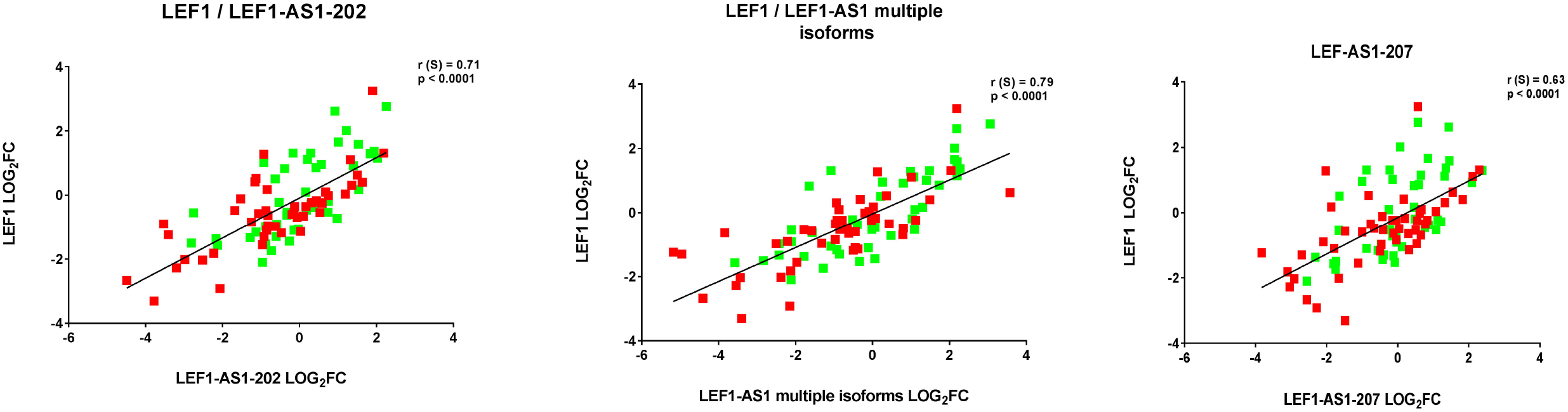
LEF1-AS1-202, LEF1-AS1 multiple isoforms and LEF1-AS1-207 positively correlate with LEF1 expression levels. Correlations for LEF1-AS1-202 (A), LEF1-AS1 multiple isoforms (B) and LEF1-AS1-207 (C) were calculated using Spearman’s correlation test. Coefficients and p values are indicated. Red and green squares indicate MPS patients (n= 48) and no MPS patients (n = 49), respectively.

As for the LEF1-AS1-202 isoform, no significant differences were observed in the expression levels of all other LEF1-AS1 isoforms and of LEF1 when patients were grouped according to MNS (Figure S4).

### 2.5 LEF1-AS1 isoforms and LEF1 correlated with relevant clinical parameters

Clinically relevant haematological tests were conducted in the peripheral blood harvested at follow-up of long COVID patients (Table S1).

Although still within reference values, patients with MPS exhibited significantly higher white blood cell levels (p <0.0007) and lower lymphocyte levels (p<0.03). Conversely, no significant differences were observed in the comparison of clinical parameters between patients with or without MNS.

Interestingly, Figure 4 shows some weak but significant correlations. All LEF1-AS1 isoforms were inversely correlated with the age of long COVID patients; moreover, LEF1-AS1 and LEF1 negatively correlated with monocyte and eosinophil levels.

**Figure 4.**
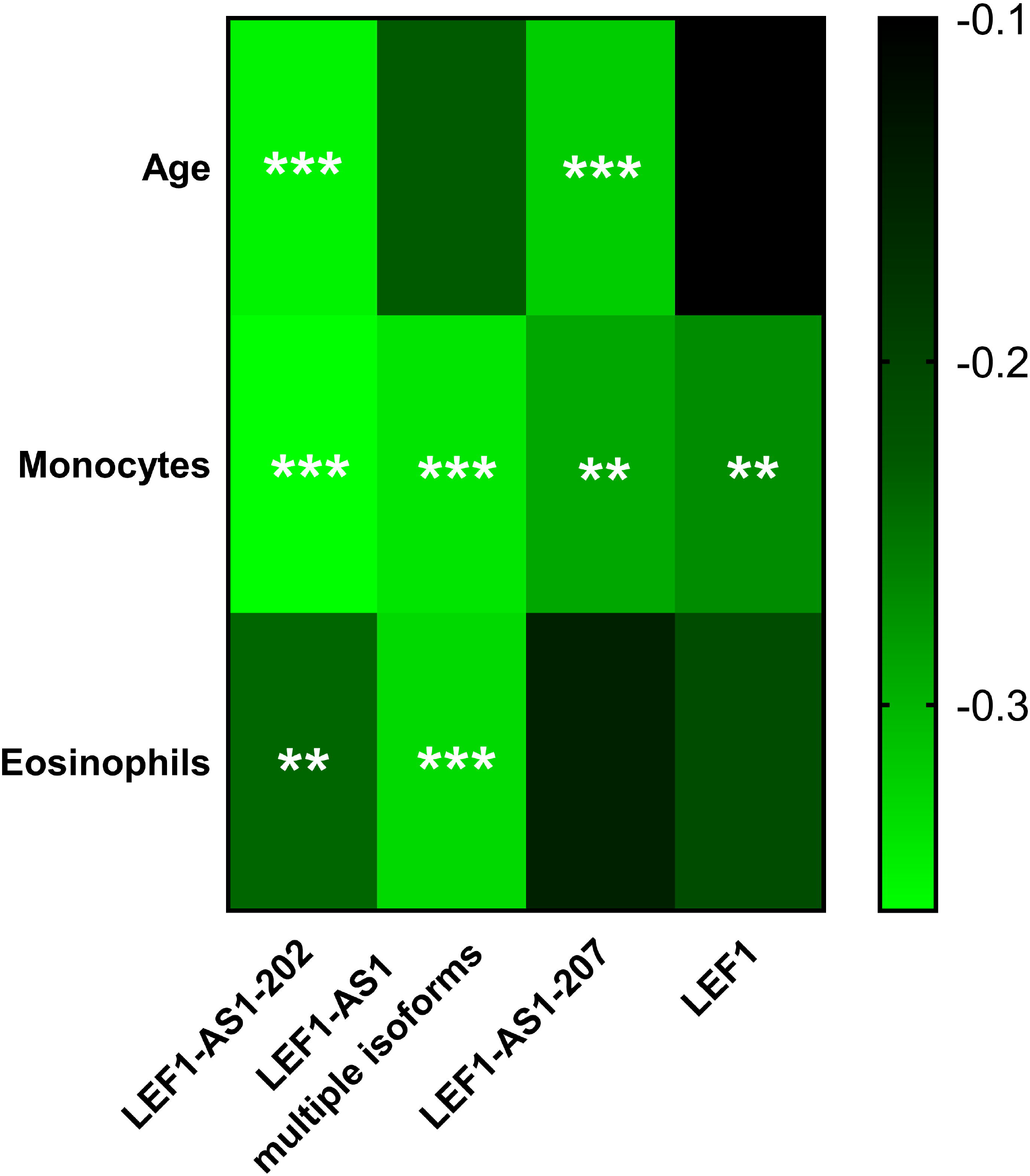
Heat-map displaying the correlations of LEF1-AS1 isoforms and LEF1 with clinically relevant parameters. Clinical parameters were measured at the moment of the interview; brighter green colour indicates a stronger negative correlation while red colour indicates a positive correlation. R coefficients were calculated with Spearman’s correlation test and stars indicate statistical significance (**p≤ 0.01;***p≤ 0.001; n= 97).

## 3. Discussion

Given the wide range of long COVID symptoms, identifying potential mechanistic elements and biomarkers for long COVID is necessary to enhance the clinical evaluation and management of affected patients. Some investigations suggest that the likelihood of experiencing long COVID symptoms decreases over time [26], but multiple studies report that the incidence of long COVID symptoms remains high even more than a year after the acute phase of COVID-19 [5,27–29]. These findings align with the symptom incidence observed in our patient cohort, in which the most common reported symptoms include cognitive issues, sensorimotor disturbances, shortness of breath, and fatigue, all of which significantly affect daily life.

In this study, we evaluated the expression of a subset of ncRNA biomarkers in long COVID patients, enrolled at a median of 18 months post-hospitalization for COVID-19. Specifically, we assessed the possible modulation of miRNA and lncRNAs previously found by our group to be deregulated in acute COVID-19 patients [23–25]. We found that LEF1AS1 levels were significantly reduced in the PBMCs of patients with MPS compared to those without MPS.

Data indicate the specificity and the potential relevance of LEF1AS1 deregulation. Indeed, while other lncRNAs, such as HCG18 or lncCEACAM21 were not modulated, all LEF1-AS1 isoforms tested were similarly deregulated in MPS patients. Moreover, when grouping patients for the persistence of MNS, no significant changes were observed: the physio-pathological reasons for this difference were not investigated, but it may indicate the implication of LF1-AS1 in MPS symptoms. Accordingly, we observed a positive correlation between LEF1-AS1 and LEF1, and an inverse correlation between LEF1-AS1 isoforms and LEF1 with relevant clinical parameters.

LEF1-AS1 expression levels showed a weak but significant negative correlation with the age of long COVID patients, in line with the evidence indicating age as a risk factor for long COVID [10,30]. Moreover, despite monocytes and eosinophils levels did not change significantly in MPS patients, they negatively correlated with LEF1-AS1 and LEF1 expression levels, possibly indicating a weak but potentially relevant pro-inflammatory condition. Accordingly, it has been shown that acute COVID-19 affects the relative proportions of monocyte subsets, leading to the activation of classical monocytes, and of pro-inflammatory cytokine production, that persists at later stages of the disease [31]. Moreover, neutrophil and eosinophil responses remain abnormal for several months after the acute COVID-19 phase [32].

LEF1-AS1 is an antisense RNA to the lymphoid enhancer binding factor 1 (LEF1) gene, which encodes a transcription factor expressed mainly, but not exclusively, in pre-B and T cells. This factor is involved in cell proliferation, activation of genes in the Wnt/β-catenin pathway, and regulation of systemic inflammation. In cancer cells, LEF1-AS1 has a pro-oncogenic role, accelerating tumorigenesis and supporting cell proliferation and invasion in glioma [33] and non-small-cell lung cancer [34]. In COVID-19, the retained intron isoform 202 of LEF1-AS1 was identified as a potential biomarker for both the outcome and severity of the disease [24,25], displaying lower levels in more severe and nonsurviving COVID-19 patients. Accordingly, neutrophil-to-lymphocyte ratios display an inverse correlation to the expression of LEF1-AS1 [24]. In long COVID patients, this inverse correlation was not observed, implicating that the decreased levels of LEF1 and LEF1-AS1 in MPS may not simply be attributed to lower levels of lymphocytes. In long COVID, the significant reduction of all LEF1-AS1 isoforms tested suggests that the entire locus of this lncRNA may be involved in the underlying pathological mechanisms of persistent MPS. Moreover, the positive correlation between all the LEF1-AS1 isoforms and LEF1 supports the idea of co-dependent regulation of these molecules, suggesting a potential common pathway influenced by long COVID.

Some studies reveal that LEF1-AS1 can interact with LEF1 through different mechanisms. In colorectal cancer, LEF1-AS1 recruits MLL1 histone methyltransferase to the promoter of LEF1, stimulates H3K4me3 methylation, and activates LEF1 transcription [35]. Unfortunately, this study did not distinguish between LEF1-AS1 isoforms: since they have significantly different sequences, studying each individually is of great importance, as they may work through different mechanisms. Indeed, Lu et al. found that LEF1-AS1-207 sponges the RNA binding protein HNRNPL which, in turn, stabilizes LEF1 mRNA [36]. In prostate cancer cells LEF1-AS1 de-represses LEF1 expression by sponging away miR-330-5p [37]. While this is very interesting, to the best of our knowledge, no evidence of miR-330-5p deregulation in COVID-19 has been reported so far, suggesting that this mechanism may not be particularly relevant in long COVID.

Our study has several limitations. First, this is a monocentric study, and our cohort only includes patients who required hospitalization during the acute phase of COVID-19. Thus, LEF1AS1 and LEF1 levels may not display long-term alterations in the PBMCs of patients that were affected by milder symptoms during acute COVID-19. Second, participants were asked to return to our hospital for the questionnaire and blood sample collection, possibly biasing the enrolment: individuals with more severe long COVID symptoms who were unable to visit the hospital independently could have been excluded. Third, physical symptoms and diseases recorded during hospitalization were obtained from patient files, providing reliable data. However, pre-existing symptoms related to MNS, such as anxiety and depression, were collected partially from patient files and partially from follow-up interviews. This latter source might be biased due to the subjective interpretation and memory of the patients. Fourth, we did not use instrumental or objective measures to validate the presence and severity of long COVID symptoms reported by patients.

In conclusion, our study identified LEF1-AS1 as a promising potential biomarker for MPS of long COVID, contributing to the understanding of the molecular mechanisms underlying the condition. While further studies are needed to validate and expand these findings, our discoveries offer new perspectives for the diagnosis and management of long COVID.

## 4. Materials and Methods

### 4.1 Ethics approval and consent to participate

The study was performed in full compliance with the Declaration of Helsinki. The experimental protocol of IRCCS Policlinico San Donato (protocol number 39/INT/2022, of April 06, 2022) was approved by the Institutional Ethics Committee of San Raffaele Hospital. Prior to enrolment, participants were requested to provide informed consent, as previously approved by the ethics committee.

### 4.2 Patient selection and sample collection

Subjects hospitalized at IRCCS Policlinico San Donato due to acute COVID-19 between January 2021 and June 2022 constituted the eligible patient population for the “long COVID” protocol. Overall, 98 patients were enrolled. Patients aged between 18 and 90 years old were contacted through a telephone call during the period between July 2022 and December 2022. The protocol of the study included extracting pertinent data about COVID-19 hospitalization, the administration of the long COVID questionnaire validated by the Italian Ministry of Health Data [5] via in-person interviews conducted by a biologist and a medical doctor, and the collection of the peripheral blood sample. Platelet-poor plasma and Peripheral Blood Mononuclear Cells (PBMC) samples were prepared according to internal Standard Operating Procedures by BioCor Biobank at IRCCS Policlinico San Donato, part of BBMRI-ERIC and BBMRI.it and operating according to national and international guidelines.

The questionnaire included items comprising a wide array of symptoms related to long COVID. For each symptom, patients were asked to specify whether it was still present at the time of the interview, had been present but resolved, or was present intermittently. Based on the symptoms not present before the hospitalization for COVID-19 and still present at the time of the interview, patients were classified in two different ways, as previously described [5]. Those who exhibited at least one persistent symptom among fever, persistent cough, chronic fatigue or shortness of breath, were classified as patients affected by major physical symptoms (MPS), while those who exhibited at least one persistent symptom among anxiety, depression, brain fog and memory dysfunction, were classified as major neuropsychological symptoms (MNS) patients. Participant data were collected at the time of the interview using the Research Electronic Data Capture (RedCAP) platform.

### 4.3 RNA isolation and RT-qPCR quantification

RNA isolation from PBMC samples was carried out using Trizol RNA Extraction reagent (#FS-881-200, Società Italiana Chimici) as previously described [38]. Before reverse transcription, 1 μg of RNA was digested with 1 U of DNase I (#18068-015, ThermoFisher Scientific) according to manufacturer’s protocol. The digestion mixture was incubated at 37°C for 30 minutes and then the reaction was inactivated by adding 50 mM EDTA and incubating the samples at 65°C for 10 minutes. Reverse transcriptase reactions were then conducted in the presence (RT+) or in the absence (RT-) of the enzyme, followed by PCR amplification. RT-samples did not show any amplification signal. lncRNA levels were measured with GoTaq® qPCR Master Mix (#A6002, Promega) and were normalized to UBC levels. Primer sequences are listed in Table S2; alignment was performed using GRCh38/hg38 Ensembl Genome Browser. The size of the LEF1-AS1 isoforms and LEF1 amplification products were visualized using 2% agarose gel electrophoresis (Figure S3 and Figure S5). RNA isolation from platelet-poor plasma samples was performed with NucleoSpin miRNA Plasma (#740981.50, MACHEREY–NAGEL) following the manufacturer’s instructions. miRNA expression levels were investigated as previously described [23]. Specific miRNA primers were provided by ThermoFisher Scientific and the expression values were normalized to U6 levels. All RNA species were quantified using CFX Opus 96 Real-time PCR System (BIO-RAD) and relative expression was calculated using the comparative Ct (Delta Delta Ct) method [39].

### 4.4 Statistical analysis

Continuous variables are shown as mean ± standard error of the mean (SEM). Group-wise comparisons were conducted using either the Mann–Whitney test or unpaired t-test, as appropriate. All statistical tests were two-sided, with a significance level set at p < 0.05. Correlation analyses were performed using Spearman’s test. All statistical analyses were performed using GraphPad Prism v.8.3.0 software (GraphPad Software Inc.).

## 5. Patents

### Supplementary Materials

The following supporting information can be downloaded at: www.mdpi.com/xxx/s1, Figure S1. No significant changes in miR-144-3p expression levels between no MPS and MPS patients and between no MNS and MNS patients; Figure S2. No significant differences in lncRNA expression levels between patients with and without MNS; Figure S3. LEF1-AS1 locus transcript annotation and qPCR primer pairs; Figure S4. No significant differences in LEF1-AS1 isoforms and LEF1 expression levels between patients with and without MNS; Table S1. Haematological tests and their respective reference values for LONG COVID patients at follow-up; Table S2. Primer sequences; Figure S5. Amplicons of LEF1 and LEF1-AS1 isoforms.

## Author Contributions

F.M., A.M., C.G., and G.S.: conceptualization and study design; A.M. and S.N.P.: performing the experiments; M.R., L.V.R., and R.C.: patient enrollment; A.M. and F.M.: writing the original draft; A.M., S.N.P., M.R., C.G., L.V.R., R.C., G.S. and F.M. writing review and editing.

## Institutional Review Board Statement

The study was conducted according to the guidelines of the Declaration of Helsinki, and approved by the Ethics Committee of Ospedale San Raffaele (protocol number 39/INT/2022, of April 06, 2022).

## Informed Consent Statement

Informed consent was obtained from all subjects involved in the study. Written informed consent has been obtained from the patients to publish this paper.

## Data availability statement

Data that support the findings of this study, not publicly available due to privacy or ethical restrictions, can be obtained on request from the corresponding author.

## Acknowledgments

We thank Ekaterina Baryshnikova, Martina Anguissola, and Sara Pugliese for patient recruitment, Arianna Pinton for sample collection, and all the clinical and technical personnel of PSD that made this study possible. During the preparation of this work the authors used ChatGPT (OpenAI, San Francisco, CA) in order to improve the grammar and clarity of the manuscript. After using this tool/service, the authors reviewed and edited the content as needed and take full responsibility for the content of the publication.

