## Supplementary figures and tables for "LEF1-AS1 deregulation in the peripheral blood of patients with persistent post-COVID symptoms"

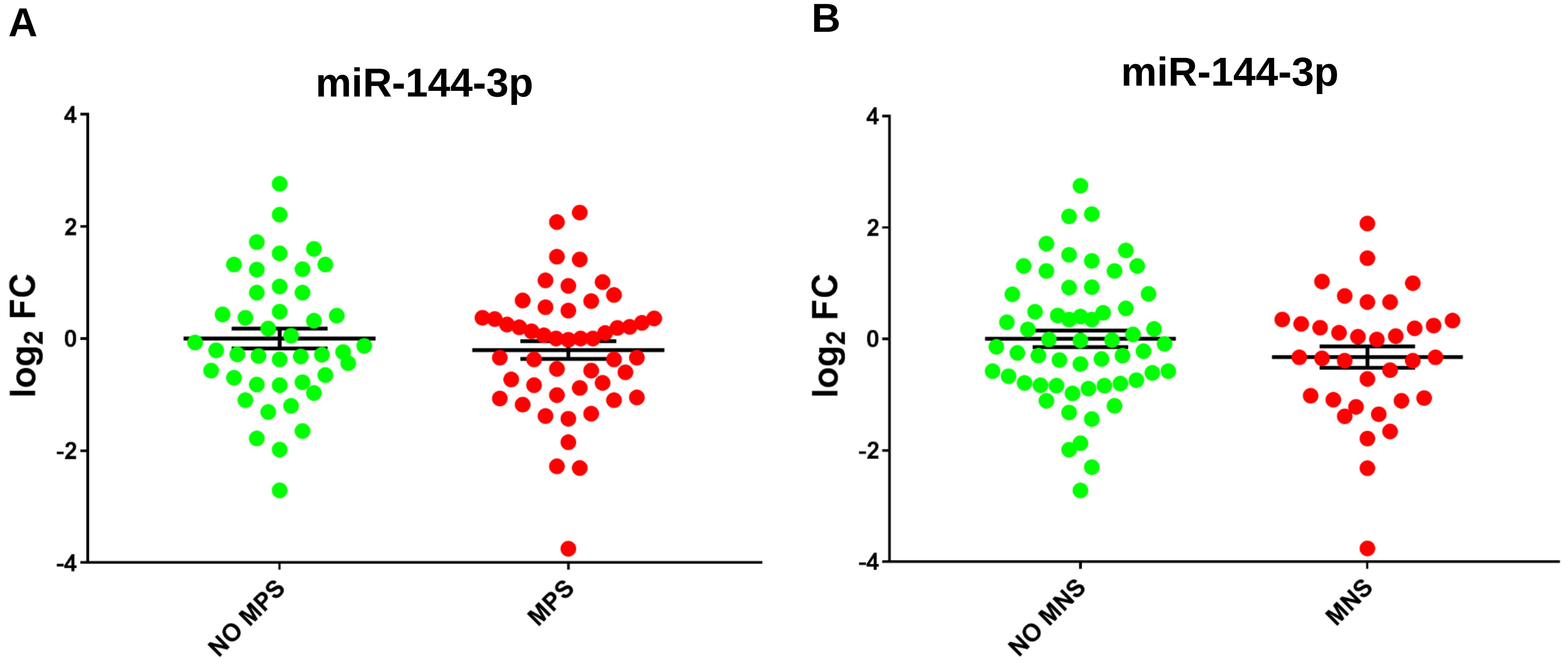


**Figure S1. No significant changes in miR-144-3p expression levels between no MPS and MPS patients and between no MNS and MNS patients.** miR-144-3p plasma levels were measured by qPCR in patients with or without MPS (A) or MNS (B). Values are expressed as $\log_2$ fold change and shown as dot-plots indicating mean ± SEM. Unpaired t-test (two groups) was used for statistical comparison: differences were not statistically significant. No MPS patients n = 43; MPS patients n = 48. No MNS patients n = 56; MNS patients n = 35.

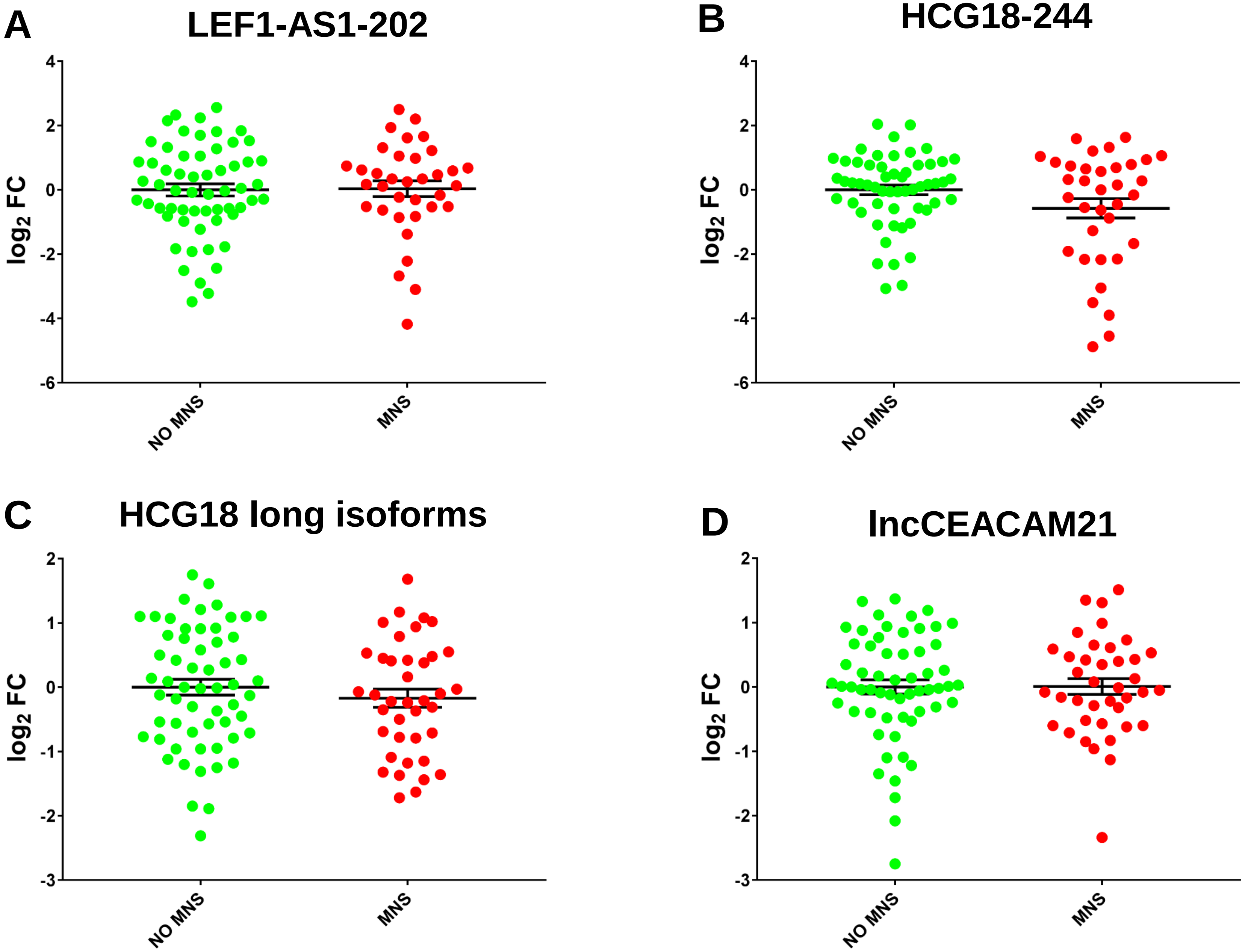


**Figure S2. No significant differences in lncRNA expression levels between patients with and without MNS.** LEF1-AS1-202 (A), HCG18-244 (B), HCG18 long isoforms (C) and lncCEACAM21 (D) were measured in total RNAs extracted from the PBMC of no MNS (n = 59) and MNS patients (n = 35-39). Values are expressed as $log_2$ fold change and shown as dot-plots, indicating mean ± SEM. Unpaired t-test (two groups) was used for statistical comparison: differences were not statistically significant.

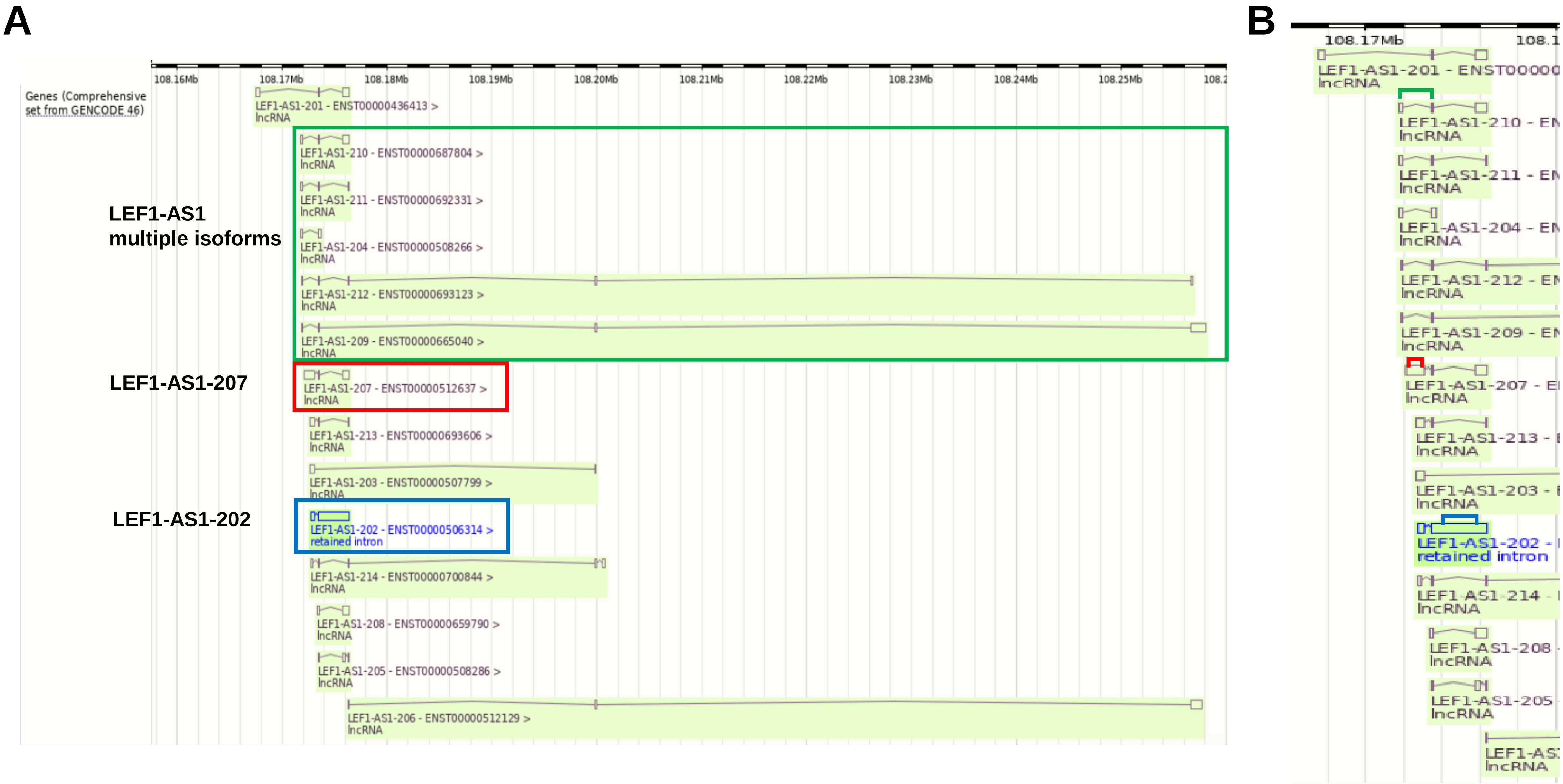


**Figure S3. LEF1-AS1 locus transcript annotation and qPCR primer pairs.** In panel A, the green box represents all isoforms identified as multiple isoforms, while the red and blue boxes represent LEF1-AS1-207 and LEF1-AS1-202 isoforms, respectively. Panel B zooms in on 5′ region, where the green, red and blue brackets indicate the position of the primers used to detect LEF1-AS1 multiple isoforms, LEF1-AS1-207 and LEF1-AS1-202, respectively. The representation of the locus was obtained by Ensembl genome browser (Chromosome 4: 108,167,525-108,258,037 forward strand; GRCh38:CM000666.2).

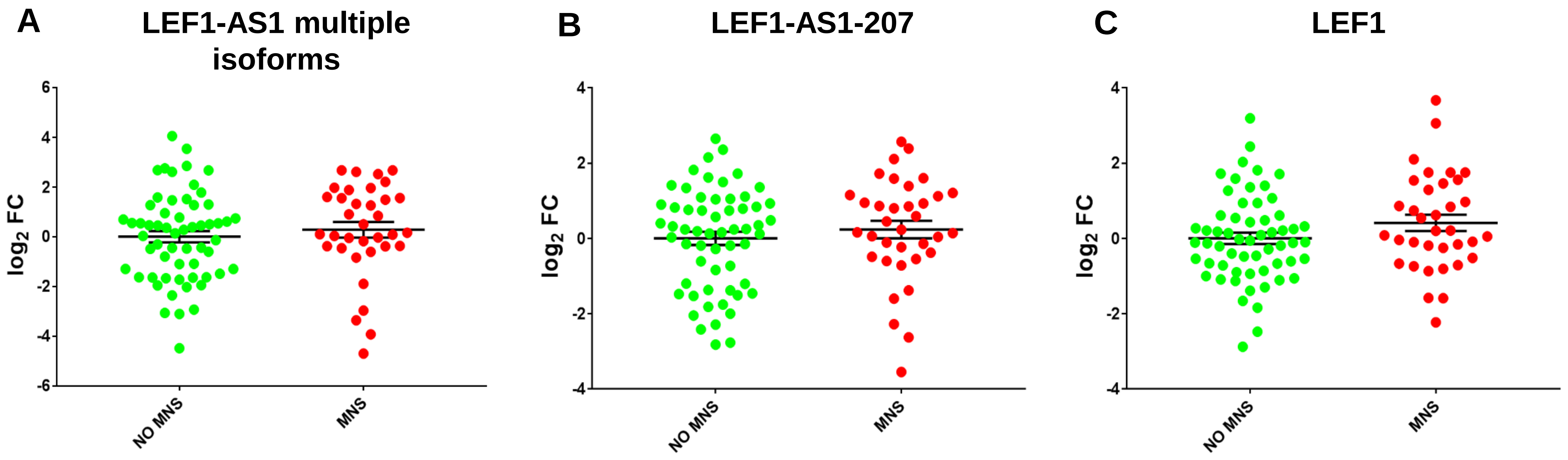


**Figure S4. No significant differences in LEF1-AS1 isoforms and LEF1 expression levels between patients with and without MNS.** LEF1-AS1 multiple isoforms (A), LEF1-AS1-207 (B), and LEF1 (C) were measured in total RNAs extracted from the PBMCs of patients with MNS (MNS, n = 39) or without MNS (no MNS, n = 59). Values are expressed as $\log_2$ fold change and shown as dot-plots indicating mean ± SEM. Unpaired t-test (two groups) was used for statistical comparison: differences were not statistically significant.

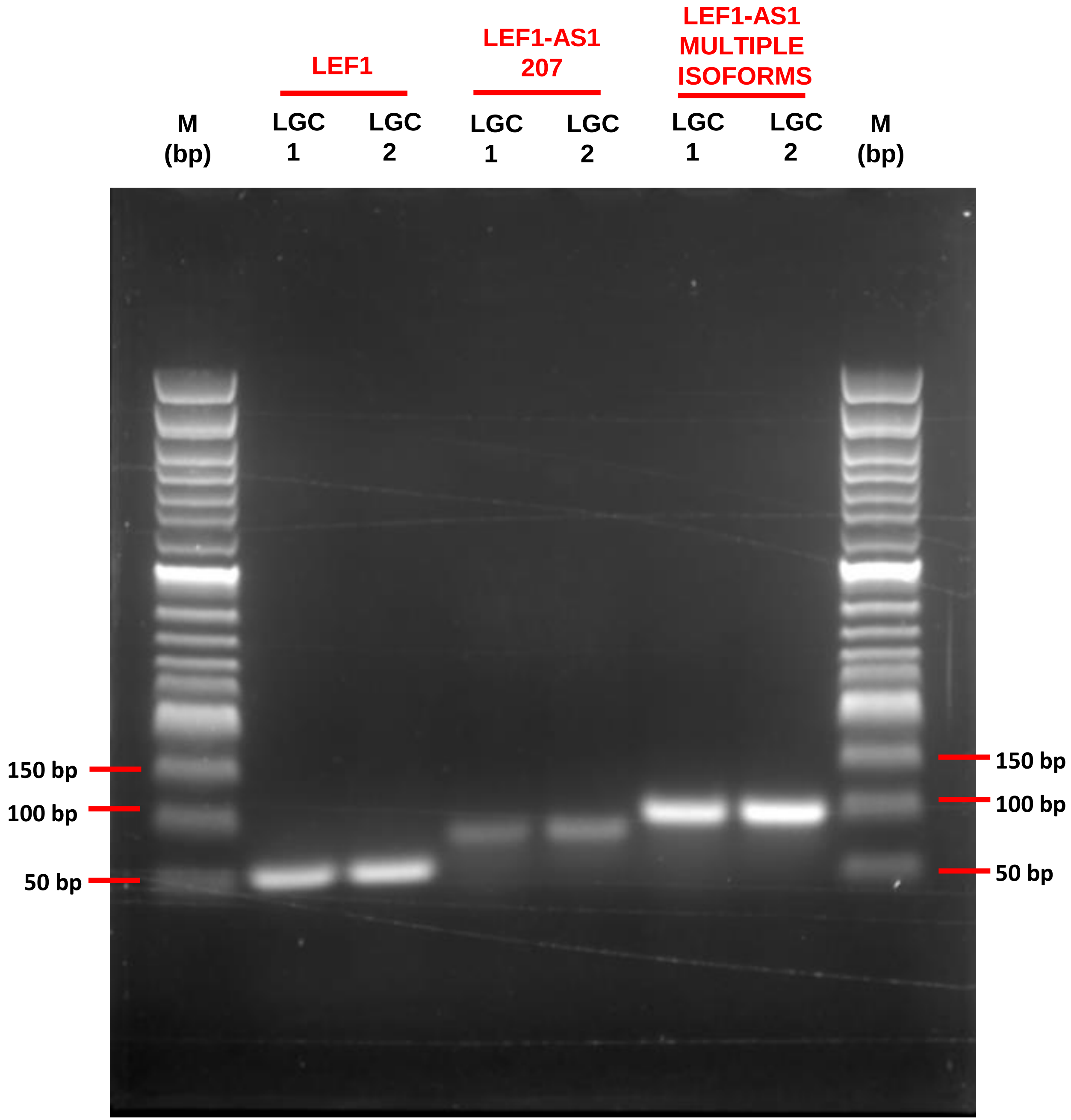


**Figure S5. Amplicons of LEF1 and LEF1-AS1 isoforms.** Total RNA was extracted from the PBMC of 2 patients of the long COVID study. DNAse digestion and reverse transcription followed by PCR amplification for 40 cycles were performed in order to detect LEF1, LEF1-AS1-207, and LEF1-AS1 multiple isoforms. The agarose electrophoretic analysis shows a single detectable amplicon for each primer pair, indicating the specificity of the qPCR reactions.

**Table S1**. Haematological tests and their respective reference values for LONG COVID patients at follow-up

| | Reference values | MPS (N = 48) | NO MPS (N = 50) | *p value* | MNS (N = 39) | NO MNS (N = 59) | *p value* |
|---|---|---|---|---|---|---|---|
| Haemoglobin | 14-18 g/dL | 14.4 ± 0.46 | 14.45 ± 1.41 | *ns* | 13.55 ± 0.45 | 15.0 ± 1.38 | *ns* |
| Haematocrit | 41-53 % | 43.7 ± 0.71 | 43.2 ± 1.16 | *ns* | 41.4 ± 0.72 | 44.20 ± 1.15 | *ns* |
| Red blood cells | 4.50-5.90 10^12/L | 4.84 ± 0.28 | 4.78 ± 0.35 | *ns* | 4.55 ± 0.31 | 4.85 ± 0.35 | *ns* |
| White blood cells | 4.00-11.30 10^9/L | 7.30 ± 0.80 | 6.40 ± 0.77 | *0.0007* | 6.95 ± 0.90 | 6.64 ± 1.04 | *ns* |
| Neutrophils | 1.90-8.00 10^9/L | 4.41 ± 0.56 | 5.03 ± 5.08 | *ns* | 3.91 ± 0.74 | 4.05 ± 0.83 | *ns* |
| Lymphocytes | 1-4.80 10^9/L | 2.06 ± 1.34 | 3.76 ± 0.75 | *0.03* | 2.45 ± 1.19 | 2.01 ± 1.01 | *ns* |
| Monocytes | 0.16-1 10^9/L | 0.43 ± 1.08 | 0.35 ± 0.19 | *ns* | 0.42 ± 0.27 | 0.41 ± 0.24 | *ns* |
| Eosinophils | 0-0.80 10^9/L | 0.16 ± 0.34 | 0.14 ± 0.5 | *ns* | 0.14 ± 0.27 | 0.16 ± 0.36 | *ns* |
| Basophils | 0-0.20 10^9/L | 0.04 ± 0.13 | 0.03 ± 0.15 | *ns* | 0.04 ± 0.13 | 0.04 ± 0.16 | *ns* |
| Platelets | 150-400 10^9/L | 221.0 ± 50.67 | 216.5 ± 58.28 | *ns* | 221.0 ± 50.22 | 215.0 ± 36.15 | *ns* |
| Total bilirubin | 0.30-1.20 mg/dL | 0.68 ± 0.31 | 0.7 ± 0.75 | *ns* | 0.56 ± 0.47 | 0.72 ± 0.67 | *ns* |
| GPT/ALT | 9-40 U/L | 24.0 ± 4.82 | 24.5 ± 3.81 | *ns* | 22.0 ± 2.54 | 28.0 ± 5.25 | *ns* |
| Creatinine | 0.60-1.10 mg/dL | 0.94 ± 0.43 | 0.86 ± 0.53 | *ns* | 0.89 ± 0.51 | 0.93 ± 0.48 | *ns* |
| Sodium | 136-145 mmol/L | 141.0 ± 0.19 | 140.0 ± 0.88 | *ns* | 140.0 ± 0.17 | 141.0 ± 0.17 | *ns* |
| Potassium | 3.50-5.10 mmol/L | 4.34 ± 0.21 | 4.31 ± 0.39 | *ns* | 4.31 ± 0.25 | 4.34 ± 0.30 | *ns* |
| LDH | 120-246 U/L | 206.0 ± 2.86 | 196.0 ± 35.62 | *ns* | 201.0 ± 27.81 | 200.5 ± 27.20 | *ns* |
| CRP | <5 mg/L | 0.28 ± 3.52 | 0.46 ± 0.86 | *ns* | 0.26 ± 2.74 | 0.47 ± 0.71 | *ns* |
| INR | | 1.09 ± 0.10 | 1.06 ± 0.36 | *ns* | 1.08 ± 0.10 | 1.07 ± 0.36 | *ns* |
| aPTT | 21.1-37.7 sec | 27.90 ± 1.04 | 28.0 ± 8.26 | *ns* | 27.25 ± 0.75 | 28.70 ± 1.50 | *ns* |
| D-dimer | <550.00 ug/L | 382.0 ± 317.9 | 269.0 ± 310.89 | *ns* | 460.0 ± 334.1 | 257.0 ± 333.25 | *ns* |
| Fibrinogen | 170-420 mg/dL | 315.0 ± 68.77 | 301 ± 61.19 | *ns* | 315.0 ± 59.7 | 305.0 ± 56.31 | *ns* |

Values are expressed as median ± SEM.
GPT/ ALT: alanine aminotransferase; LDH: Lactate dehydrogenase; CRP: c-reactive protein; INR: international normalised ratio; aPTT: Activated Partial Thromboplastin Time.

**Table S2**. Primer sequences

| Table S1: primer sequences | |
|---|---|
| **Gene** | **Primer pairs** |
| **HCG18-244** | Forward: GTGGGCTATATTAAGTGGGAAT<br>Reverse: TCTTCTCTATATCTCAAGGTATGC |
| **HCG18 long isoforms** | Forward: AGGTACTGGTGGTGTGGCTA<br>Reverse: GGTCATCATCAGTGGGGCTC |
| **lncCEACAM21** | Forward: CAAGTATCTCTTCTCAACGGAAA<br>Reverse: ATCATTATTCACGCATCCACAA |
| **LEF1-AS1-202** | Forward: GTCCATGCTATGACCATCTCCA<br>Reverse: ACACGAGTTAAGGCACATTCAC |
| **LEF1-AS1 multiple isoforms** | Forward: CACAAGAACAGAGGAGCGGG<br>Reverse: AGCACATCCCCCAACATTCT |
| **LEF1-AS1-207** | Forward: AAGGACGAGAGAAAAGCAC<br>Reverse: CACACAAAGGGGAAGACC |
| **LEF1** | Forward: AGAACACCCCGATGACGGA<br>Reverse: GAGGGTCCCTTGTTGTAGAGG |
| **UBC** | Forward: GATCGCTGTGATCGTCACTTGACAA<br>Reverse: AGTCAGACAGGGTGCGCCCA |
